# No isolate, no problem: Using a novel insertion sequence PCR to link rats to human shigellosis cases in an underserved urban community

**DOI:** 10.1101/2023.03.02.530678

**Authors:** Gordon Ritchie, Victor Leung, Chelsea G. Himsworth, Kaylee A. Byers, Lisa K.F. Lee, Samuel D. Chorlton, Aleksandra Stefanovic, Marc G. Romney, Nancy Matic, Christopher F. Lowe

## Abstract

**Introduction:** During an investigation into a cluster of *Shigella flexneri* serotype 2a cases in an underserved community, we assessed the relatedness of human and rat *S. flexneri* isolates utilizing a novel PCR targeting insertion sites (IS-PCR) of mobile elements in the *Shigella* genome characteristic of the cluster strain.

**Methods:** Whole genome sequences of *S. flexneri* (n=50) associated with the cluster were analyzed. *de novo* genome assemblies were analyzed by a Geneious V10.2.6 motif search, and 2 unique IS were identified in all human *Shigella* sequences of the local cluster. Hydrolysis probe PCR assays were designed to detect these sequences consisting of forward and reverse primers to amplify across each insertion site, and a hydrolysis probe spanning the insertion site. IS-PCR was performed for three *Shigella* PCR-positive culture-negative rat intestine specimens from this community.

**Results:** Both insertion sites were detected in the *de novo* genome assemblies of all clinical *S. flexneri* isolates (n=50). Two of the three PCR-positive culture-negative rat samples were positive for both unique IS identified in the human *S. flexneri* isolates, suggesting that the rat *Shigella* spp. strains were closely related to the human strains in the cluster. The cycle threshold (Ct) values were >35, indicating that the bacterial load was very low in the rat samples.

**Conclusions:** Two unique IS were identified in clinical isolates from a community *S. flexneri* cluster. Both IS targets were identified in PCR-positive (*Shigella* spp.), culture-negative rat tissue and clinical isolates from humans, indicating relatedness.

## Introduction

*Shigella* spp. have high potential for transmission from person-to-person through fecal-oral contact due to their low infectious dose (10-100 organisms)(1). Humans are thought to be the only natural reservoir for *Shigella* spp., and outbreaks/clusters have been reported in a variety of settings, including daycares/schools, men who have sex with men, foodborne transmission, travelling and regions with poor hygiene and sanitation(2, 3).

There are numerous methods available for outbreak analysis (pulsed-field gel electrophoresis [PFGE], amplified fragment length polymorphism [AFLP], random amplified polymorphic DNA [RAPD], variable number tandem repeat [VNTR], multi-locus sequence typing [MLST], and core genome MLST [cgMLST]), but these methods are dependent on growth of the bacterial isolate in culture(4). Culture-dependent methods are challenging when working with fastidious organisms or samples with low burdens of pathogen. Cases of culture-negative shigellosis have been increasingly recognized due to the use of molecular diagnostics(5, 6), rendering many of these typing methods ineffective. These methods are also resource-intensive requiring specialized instrumentation and technical/bioinformatics expertise. Culture-independent typing methods that are scalable and simple to perform are currently needed to enhance outbreak investigations. Specifically for *Shigella* spp. which contain a high number of mobile genetic elements and genomic rearrangements, strain relatedness can potentially be analyzed through targeting of insertion sequences(3, 7).

In 2021, a cluster of *Shigella flexneri* serotype 2a cases occurred in an underserved neighbourhood in Vancouver, British Columbia. A specific source (e.g. housing, food, sexual exposures, travel) was not identified, but poor sanitary conditions in this community were presumed to have contributed to ongoing transmission (http://www.vch.ca/Documents/26%20Feb%202021%20Physicians%20Update%20Shigella%20Outbreak%20Hep%20A%20Cases.pdf). Rats were hypothesized to be one potential vector for ongoing transmission based on previous observations(8). Although samples from rats (*Rattus norvegicus*) recovered from this community were culture-negative for *Shigella* spp., three rat samples tested positive for *Shigella*/Enteroinvasive *E. coli* (EIEC) using polymerase chain reaction (PCR) [Biofire® FilmArray® Gastrointestinal Panel](C. Himsworth, unpub. data). We utilized a novel PCR targeting insertion sites (IS-PCR) of mobile elements in the *Shigella* genome characteristic of the outbreak strain for outbreak analysis.

## Methods

*S. flexneri* (n=50) isolates from the outbreak underwent Whole Genome Sequencing (WGS) using the GridION. All genomes were serotype 2a (*in silico* serotyping with ShigEiFinder) and MLST 245(9). Human *Shigella flexneri* isolates associated with the community outbreak were grown overnight in Mueller Hinton broth, then inactivated at 99°C for 10mins. 250µL bacterial Lysis Buffer (Roche) was added to 250µL bacterial suspension, followed by the addition of 25µL of Proteinase K (Sigma). The suspension was incubated at 65°C for 1 hour, then DNA was extracted on the MagNA Pure 24 (Roche Molecular Systems). The DNA was prepared for sequencing with the SQK-LSK109 Ligation Sequencing Kit with NBD104/114 barcoding. The DNA library was run on GridIon (Oxford Nanopore Technologies) with R9 flowcells. Raw sequencing data was basecalled with Guppy (v4.1.0) and uploaded to BugSeq for further automated analysis(10). Reads were assembled with metaFlye and polished with Racon, Medaka (https://github.com/nanoporetech/medaka) and Homopolish(11–13). A consensus FASTA sequence was constructed using bugseq.com. 109 copies of the IS1 mobile element were found in the resulting *de novo* genome assemblies by a Geneious V10.2.6 motif search. Sequences spanning these insertion sites were scanned by BLAST searches of Genbank (performed August 2022 – at the time, >10,000 *S. flexneri* and >170,000 *E. coli* genome sequences). One unique insertion site was present in all human *Shigella* sequences in the local outbreak, but not present in any other *E. coli/Shigella* sequences in Genbank (ie all publicly available *E. coli/Shigella* sequences) at the time of primer design. Another rare insertion site was identified that was present in only 40 *S. flexneri* sequences within Genbank. This site was a result of a fusion of the *acrA*-like gene with the *yegE* gene interrupted by an ISSfl2 insertion sequence. Neither IS was identified in any other ST245 strain in Genbank. A BLAST search for other gastrointestinal pathogens was conducted, but did not identify any matches with the IS-PCR products.

Rats were collected from March 23 to April 9, 2021 from six city blocks in Vancouver’s urban Downtown Eastside Neighbourhoods. City blocks were selected with highest concentration of positive human *S. flexneri* cases. We used lethal Snap-e rat traps (Kness Manufacturing Co., Iowa, United States) inserted inside of tempered PROTECTA EVO Express Bait Stations (Bell Laboratories, Wisconsin, United States). Six traps were placed in the alleyway of each city block for a total of 36 active traps. We checked traps each morning, and for trapped rats we recorded capture date and location. Following capture, rats were stored at -20°C prior to undergoing a full necropsy. To collect samples for *Shigella* testing, we collected each rat’s whole intestine. This was stored at -80°C prior to *Shigella* spp. testing. Ten rat specimens (intestine) from this community were recovered for IS-PCR (three PCR-positive culture-negative). The rat intestine samples were suspended in 700uL of phosphate buffered saline (PBS) and bead lysed with 0.2mm beads on the Qiagen TissueLyser LT for 3 minutes, then extracted on the Roche MagNA pure compact and eluted in 50uL. To increase the specificity of the PCR, hydrolysis probe PCR assays were designed to detect these sequences consisting of forward and reverse primers to amplify across each insertion site, and a hydrolysis probe spanning the insertion site (Table 1). PCR was performed on the Lightcycler® 480 (Roche Diagnostics, Laval, QC) using the IDT Gene Expression Master Mix with 0.2uM primer, 0.1uM probe, and 1mg/mL BSA. The cycling conditions were 95°C for 10mins, then 60 cycles of 97°C for 5s, 66°C stepping down to 60°C at 0.2°C /cycle for 20s, 72°C for 2s, with fluorescence determination at 510nm. All primers and probes were synthesized by Integrated DNA Technologies. PCR positive control was a 1/1000 dilution of one of the human outbreak isolates. PCR negative control was PBS that was processed at the same time as the rat samples including bead lysing and extraction. Bioanalyzer 2100 gel electrophoresis (Agilent Technologies, CA) was performed on the PCR products. PCR products were barcoded, normalized and applied to the MinION (R9.4 FLO-MIN106D) according to the Oxford Nanopore 1D Native barcoding protocol (version: NBE_9065_v109_revE_23May2018). All DNA clean-ups were performed with 2X SPRI Agencourt beads to ensure adequate DNA recovery from the low MW PCR products. Consensus sequence reads were analyzed with Geneious 10.2.6 software.

**Table 1.**
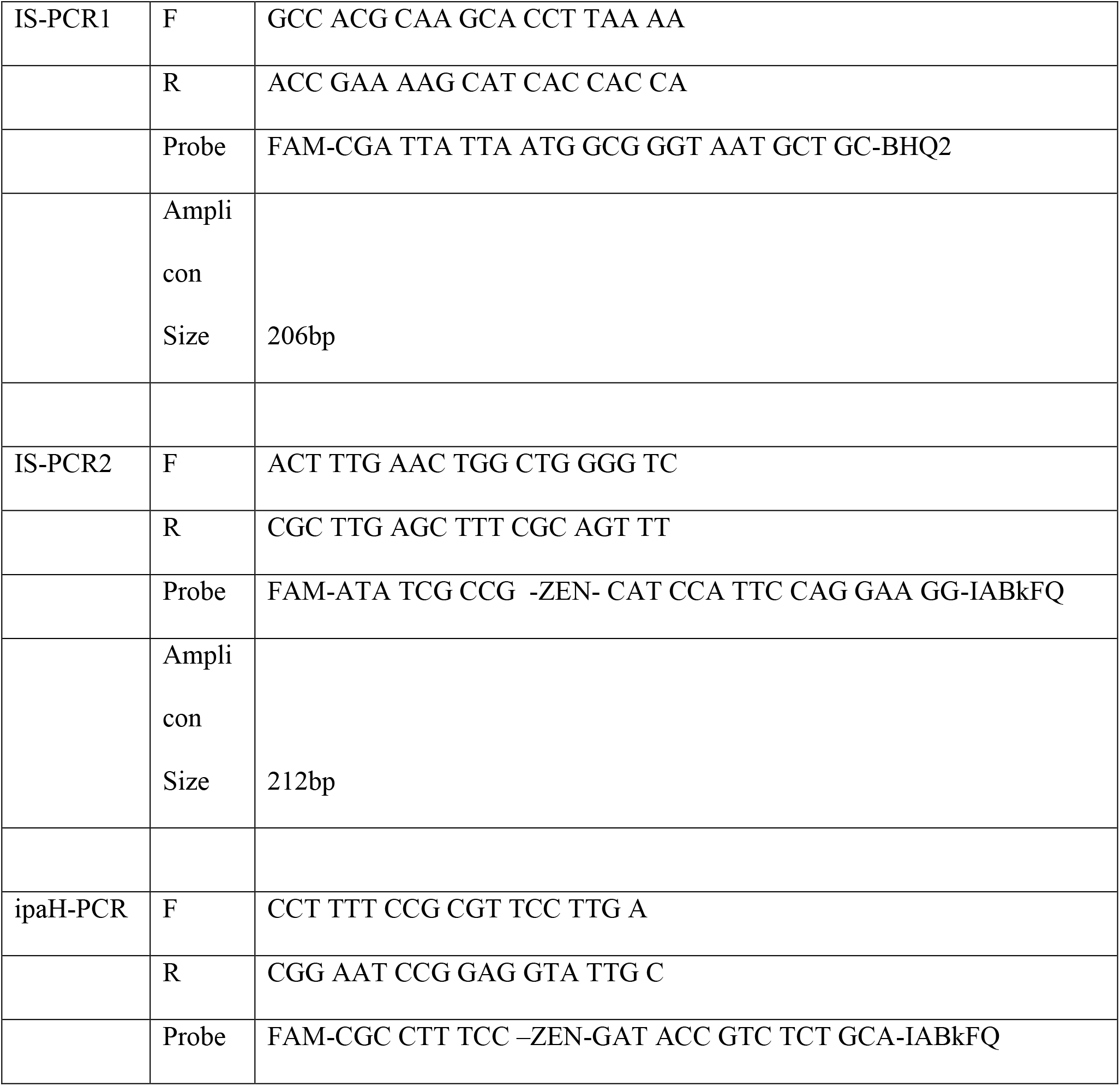
Primer and probe sequences for the Insertion Sequence PCR (IS-PCR) and *ipaH* PCR

This study was approved by the University of British Columbia’s Animal Care Committee (A20-0212) and adhered to guidelines outlined by the Canadian Council on Animal Care.

## Results

Both insertion sites were detected in the *de novo* genome assemblies of all clinical *S. flexneri* isolates (n=50), which represented samples from the course of the outbreak.

Two of the three PCR-positive culture-negative rat samples were positive for both unique IS identified in the human *S. flexneri* isolates, suggesting that the rat *Shigella* spp. strains were closely related to the human strains in the outbreak. The cycle threshold (Ct) values for the PCR were >35, indicating that the bacterial load was very low in the rat samples (Table 2). The 7 PCR-negative culture-negative rat samples were negative for both unique IS.

**Table 2.** Cycle threshold (Ct) values from direct amplification of rat intestines associated with a community *Shigella flexneri* outbreak for *ipaH* and two unique insertion sequences (IS-PCR1 and IS-PCR2)

| Rat | *ipaH* | IS-PCR1 | IS-PCR2 |
| --- | --- | --- | --- |
| 1 | 33 | 35 | 41 |
| 2 | 36 | Negative | Negative |
| 3 | 29 | 35 | 46 |

Agilent gel of IS-PCR products gave the expected amplicon sizes from the WGS of the human *S. flexneri* isolates – 206bp and 212bp. Consensus sequence reads from the positive rat samples were 100% identical to the human isolate sequences, confirming the presence of the IS in the rat samples.

## Discussion

We used a novel strain typing method (IS-PCR) to show that *Shigella spp*. from humans and rats in the same community were related. In *Shigella* spp., much of the strain-to-strain differences are due to transposase activity within mobile elements, which result in genome rearrangement. This method uses the insertion site of mobile elements in the *Shigella* genome to link bacterial strains, as insertion sites are generally stable through the course of an outbreak(3). Two mobile element insertion sites were found in all the human *Shigella* outbreak isolates that were rare in Genbank, and also detected in the rat samples by amplifying across the insertion site, with enhanced discriminatory power via a hydrolysis probe spanning the insertion site. In addition, gel electrophoresis showed identical sized amplicons for the rat samples and the human isolates, and PCR products were sequenced and matched the targeted IS. The discovery of identical rare insertion sites for IS1 in the human isolates and the rat samples demonstrates that the *Shigella* strains are related. Although IS-PCR can infer relatedness, it cannot suggest directionality of transmission. In addition, as rats are coprophagic, detection of low level *S. flexneri* DNA in rat intestines may be secondary to contact or consumption of *Shigella* contaminated food or objects rather than *Shigella* infection in the rats. Despite this, identification of the human strain in rats can provide Public Health with a potential source to target in mitigating ongoing transmission, though further research is required to understand the role rats may play in the spread of human-associated pathogens such as *Shigella*, or conversely, transmission of *Shigella* from humans to rats.

The utilization of IS-PCR enabled typing of culture-negative rat samples. There is a need for culture-independent modalities to support outbreak investigations. Stool cultures may not always be able to isolate *Shigella* spp. due to antibiotic treatment or non-viable bacteria. With the evolving use of molecular testing for the detection of gastrointestinal pathogens, an isolate may not be readily available for culture-dependent typing methods. PCR-based methods enable typing of samples with low organism burden, which may be below the limit of detection for culture or metagenomics(14). This methodology may also be amenable to outbreak/cluster investigations for other microorganisms, such as *Mycobacteria* spp., *Corynebacterium* spp. or *Staphylococcus* spp., where strain diversity is similarly achieved through mobile elements traversing throughout the genome(15).

Unique to this study was the detection of *Shigella* spp. directly from rat intestine, and evidence that it shared the same two IS sites as the human outbreak strains. A PCR-based approach was able to identify the IS sites at low levels from non-sterile tissue. This may not be possible with metagenomic sequencing as other more abundant gastrointestinal flora would be preferentially sequenced, precluding recovery of complete *Shigella* spp. genomes.

This study is limited by sample size. We applied this novel IS-PCR to only one known outbreak to date, and a potential limitation of the IS-PCR method would be the ability to detect unique insertion sites for each outbreak, which would require WGS of the index outbreak strains. A combination of IS-PCRs for rare insertion sites should also be considered, where the presence of a particular combination of 2-3 insertion sites may be uniquely identified per outbreak with good discriminatory power. Further work is also needed to assess the specificity of the IS-PCR, including testing of historical samples within our geographical region. As IS are transposable elements, there is a possibility for movement of the IS over time. For acute outbreaks over a defined time, it would be unlikely to affect the IS-PCR design, but reaffirms the need to reassess the appropriateness of the IS-PCR for subsequent outbreaks.

In conclusion, IS-PCR is a novel rapid method for determining relatedness of *Shigella* strains, which we utilized to link clinical human cases to rats. As a PCR-based approach, it may be better suited for direct PCR on tissue or samples where bacterial culture is not possible, and can serve as a complimentary modality to existing laboratory methods such as WGS for cluster/outbreak investigation.

## Acknowledgements

We thank Harveen Atwal, University of Saskatchewan and Gustavo Gabaldon, Orkin Canada for assistance with rat trapping.

